# ERK2 phosphorylates the epigenetic regulator CXXC-finger protein 1 (CFP1)

**DOI:** 10.1101/562173

**Authors:** Aroon S. Karra, Aileen M. Klein, Svetlana Earnest, Steve Stippec, Chonlarat Wichaidit, Kathleen McGlynn, David C. Trudgian, Hamid Mirzaei, Melanie H. Cobb

## Abstract

**Background:** The Ras-Raf-MEK-ERK signaling pathway is essential for proper development and homeostatic regulation in eukaryotic cells and underlies progression of several types of cancer. Many pathway functions are performed by extracellular signal-regulated kinase (ERK)1 and 2 (ERK1/2), serine/threonine protein kinases of the mitogen-activated protein kinase (MAPK) family that interact with a large number of substrates and are highly active in the nucleus.

**Results:** We identified the epigenetic regulator CXXC-finger protein 1 (CFP1) as a protein that interacts with ERK2 on chromatin. CFP1 is involved in multiple aspects of chromatin regulation, including histone methylation and DNA methylation. Here, we demonstrate the overlapping roles for ERK1/2 and CFP1 in regulation of immediate early gene (IEG) induction. Our work suggests multiple modes of co-regulation and demonstrates that CFP1 is required for an optimal signal-dependent response. We also show that CFP1 is an ERK2 substrate *in vitro* and identify several phosphorylation sites. Furthermore, we provide evidence that Su(var)3-9, Enhancer-of-zeste and Trithorax (Set)1b, a CFP1-interacting histone methylase, is phosphorylated by ERK2 and is regulated by CFP1.

**Conclusion:** Our work highlights ERK1/2 interactions with chromatin regulators that contribute to MAPK signaling diversity in the nucleus.

## Background

The extracellular signal-regulated kinases 1 and 2 (ERK1/2) function as key relays to couple extracellular signals with appropriate intracellular responses [1]. These important signaling molecules are critical for large-scale cellular processes such as proliferation, differentiation, and programmed cell death, and as such have been heavily scrutinized for their varied roles in disease progression. The physiological and pathological consequences of ERK1/2 phosphorylation on many individual substrates in several cell compartments have been documented, highlighting their far-reaching cellular functions and importance for homeostatic control [2-5].

ERK1/2 modulate gene expression through extensive interactions in the nucleus [6], where chromatin modifying enzymes regulate access to genetic information. Across species ranging from yeast to humans, trimethylation of histone H3 lysine 4 (H3K4me3) is strongly associated with promoters of actively transcribed genes [7-10]. Several studies have shown that H3K4me3 is induced by various stimuli to support transcriptional responses. Among these, MAPKs have been implicated in dynamic signal-driven histone modification, including deposition of H3K4me3, in several cell systems [11-14].

CXXC-finger protein 1 (CFP1) is a conserved component of the SETD1A and SETD1B methylation complexes and is required for maintaining specific H3K4me3 patterns in development [15-17]. CFP1 specifically binds to unmethylated DNA and can recognize unmodified di-nucleotides even in methylation-rich environments [18, 19]. CFP1 also interacts with DNA methyltransferase 1 (DNMT1), the epigenetic factor primarily responsible for maintenance of DNA 5-methylcytosine patterns established by the *de novo* methyltransferases DNMT3a and DNMT3b [20]. DNMT1 is the most abundant DNA methyltransferase in adult cells [21] and displays a distinct preference for associating with hemimethylated DNA over hypo-or hypermethylated DNA. Significantly, deletion of CFP1 in mouse embryonic stem cells (mESCs) results in a dramatic depletion of DNMT1 protein and resultant genomic DNA methylation [25].

Consistent with the conserved and multifaceted relationship between MAPK signaling and chromatin regulation, ERK1/2 are known to regulate DNMT1 [22-24]. However, a mechanistic rationale for interaction between SETD1A/B, CFP1, and DNMT1 has yet to be uncovered. DNMT1 and CFP1 contain redundant DNA targeting domains [26, 27], but unlike SETD1A/B, a dependence on CFP1 for genomic targeting of DNMT1 has not been directly demonstrated. Moreover, very little is understood about how epigenetic inputs from DNA methylation and histone methylation are integrated into gene expression outcomes in a signal-dependent manner. One likely possibility is that interactions are restricted to a subset of targeted genes or responsive to specific signaling pathways. To gain insight as to how ERK1/2 influence gene expression, we investigated ERK2 interactions with factors involved in chromatin regulation and identified a connection to CFP1. ERK1/2 and CFP1 are both known to play extensive roles in development, but less is known about CFP1 in the regulation of acute responses to extracellular cues [18, 28]. Here, we show that CFP1 is required for signal-dependent expression of ERK1/2 target genes and establish another functional relationship between MAPK signaling and chromatin dynamics.

## Results

### ERK1/2 and CFP1 interact in cells

In an effort to discover novel ERK2 substrates and binding partners, we employed a yeast-two hybrid screen designed to identify interacting factors based on the kinase activation state of ERK2. Two yeast strains were assayed in parallel, one bearing a construct expressing ERK2 alone, and another with ERK2 co-expressed with a constitutively active form of the upstream activating kinase, MEK1, termed MEK1R4F [34, 38]. A mouse neonatal cDNA library was used as prey, and among putative ERK2-interacting proteins a fragment of CFP1 encompassing the C-terminal residues 373-660 was identified with an interaction score indicating a preference for activated ERK2 (pERK2) (**Fig 1A and 4E**).

**Figure 1.**
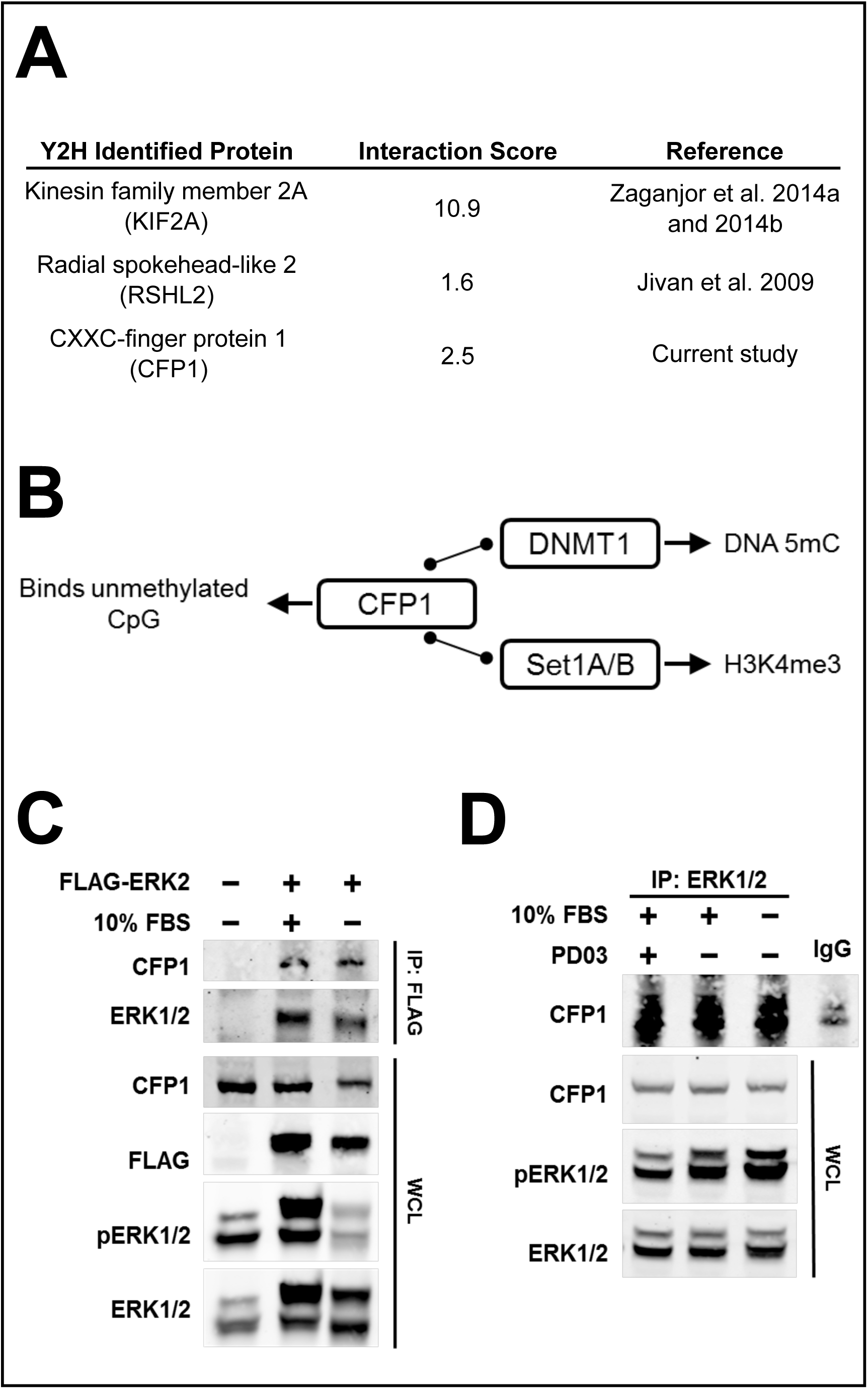
ERK1/2 interact with CFP1 in cells. **A** Summary of yeast two-hybrid results: liquid β-galactosidase assays of two-hybrid interactions between prey (neonatal mouse cDNA library) expressed as (pERK2_OD420_)/(ERK2_OD420_). **B** CFP1 interacts with several major epigenetic regulators: DNMT1 and SETD1A/B. **C** HeLa cells were transfected for 48 hours with either Flag-ERK2 or an empty vector control then starved in serum-free medium for 3 hr prior to stimulation with 10% FBS for 15 min. Isolated chromatin was subjected to limited digestion with micrococcal nuclease prior to immunoprecipitation by a monoclonal antibody directed at the FLAG epitope prior to SDS-PAGE and immunoblotting with the indicated antibodies. Whole cell lysates were loaded as controls. Results are representative of 3 independent experiments. **D** HeLa cells were treated and processed as in C. Mononucleosomes were subjected to immunoprecipitation with either an antibody recognizing endogenous ERK1/2 or rabbit IgG and immunoblotted with the indicated antibodies. Results are representative of 2 independent experiments.

To probe the interaction between CFP1 and ERK2, GST-tagged human CFP1 and fragments were purified and co-incubated with recombinant ERK1 and ERK2 activated by MEK1R4F. Although direct binding *in vitro* or in whole cell lysates was not observed, we did detect an enrichment of both ERK1/2 and CFP1 in chromatin fractions from HeLa cells (**Fig S1**). Because ERK1/2 and CFP1 are known to interact with critical chromatin modifying complexes ([39] and **Fig. 1B**), we isolated mononucleosomes from HeLa cells expressing Flag-ERK2, and performed immunoprecipitation with a resin-conjugated monoclonal Flag antibody. In addition to validating the interaction between CFP1 and ERK2 in human cells, we also included a treatment condition with PD0325901 (referred to as PD03 in the figure), a potent MEK1/2 inhibitor that effectively blocks ERK1/2 phosphorylation. Blotting for endogenous CFP1 showed that ERK2 associates with CFP1 on chromatin, regardless of the state of ERK2 activation (**Fig. 1C**). Subsequent immunoprecipitation experiments with antibodies that directly target ERK1/2 indicate this interaction is maintained by endogenous ERK2 and CFP1 on mononucleosomes (**Fig. 1D**).

### Depletion of CFP1 or inhibition of ERK1/2 signaling blunts serum-responsive gene induction

Due to the known roles of CFP1 in regulation of DNA methylation (5mC) and histone H3K4me3 deposition we investigated the effects of either CFP1 depletion or inhibition of ERK1/2 signaling on global levels of these epigenetic marks. Examination of histones acid-extracted from nuclear fractions indicated H3K4me3 is perturbed neither upon knockdown of CFP1 (**Fig. 2A**) nor inhibition of ERK1/2 signaling (**Fig. 2B**). Similarly, genomic 5mC content of cells treated with siRNA directed at CFP1 or the MEK1/2 inhibitor PD0325901 was not altered compared to control treatments (**Fig. 2C and 2D**, respectively).

**Figure 2.**
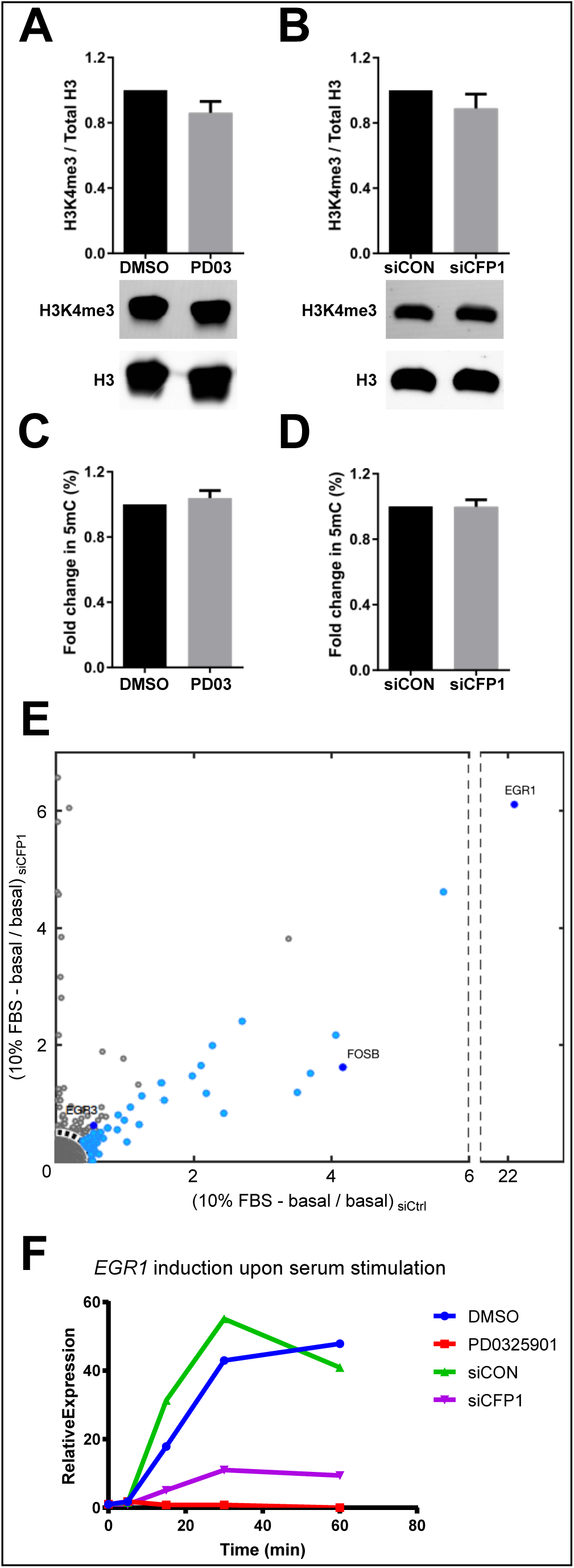
ERK1/2 and CFP1 are required for optimal serum response. **A** Genomic 5mC content was measured by ELISA for HeLa cells treated with siRNA directed at CFP1 (gene name *CXXC1*) for 72 hr. Results are representative of 3 independent experiments. **B** Global 5mC content was measured for HeLa cells treated with 500 nM PD0325901 (MEK1/2 inhibitor) or DMSO for 24 hr as in A. **C** Acid-extracted histones from HeLa cells treated as in A. and immunoblotted for total histone H3 and H3K4me3. Results from 3 independent experiments are expressed as H3K4me3 signal relative to total histone H3. **D** Acid-extracted histones from HeLa cells treated with 500 nM PD0325901 (MEK1/2 inhibitor) or DMSO for 24 hr were measured for H3K4me3 as in C. **E** Microarray analysis performed on RNA extracted from HeLa cells subjected to 72 hr of siRNA treatment with either control or CFP1-targeted oligonucleotides after serum depletion and stimulation. **F** *EGR1* mRNA induction measured by qPCR upon MEK1/2 inhibition or CFP1 depletion. Results are representative of 2 independent experiments.

CFP1 has also been shown to regulate rapid target gene-induction in B-cells [18], prompting us to test whether CFP1 is required for an ERK1/2-mediated immediate early gene (IEG) response in HeLa cells. Indeed, microarray analysis of RNA purified from cells with depleted CFP1 under serum-starved and serum-stimulated conditions revealed blunted expression of several genes, including *EGR1* (**Fig. 2E**). An extended time course further revealed that cells depleted of CFP1 or treated with PD0325901 displayed markedly similar inabilities to activate *EGR1* (**Fig. 2F**). In some other cases, serum-responsive genes did not demonstrate mutually overlapping induction profiles when comparing depletion of CFP1 and ERK1/2 signaling inhibition (**Fig. 3**). *KLF10*, for instance, is hampered under both basal and stimulated conditions when CFP1 is depleted (**Fig. 3A**). Similarly, *DUSP1* induction is exacerbated upon both CFP1 knockdown and ERK1/2 signaling inhibition (**Fig. 3B**). Interestingly, while basal amounts of *DUSP5* transcript did not change significantly after 72 hours (**Fig. S3**), starvation followed by stimulation with FBS revealed blunted gene induction under both siCFP1-and PD0325901-treated conditions (**Fig. 3C**). The prototypical IEGs *EGR3 and FOSB*, much like *DUSP5*, were also compromised by both treatments, albeit to differing degrees (**Fig. 3D and 3E**, respectively), behaving more like *EGR1*. Interestingly, Diverse signaling of ERK1/2 and CFP1 at target genes was not entirely unexpected and suggests multiple modes of co-regulation (**Fig. 3F**). These modes of regulation are not mutually exclusive and are likely dependent on specific signals.

**Figure 3.**
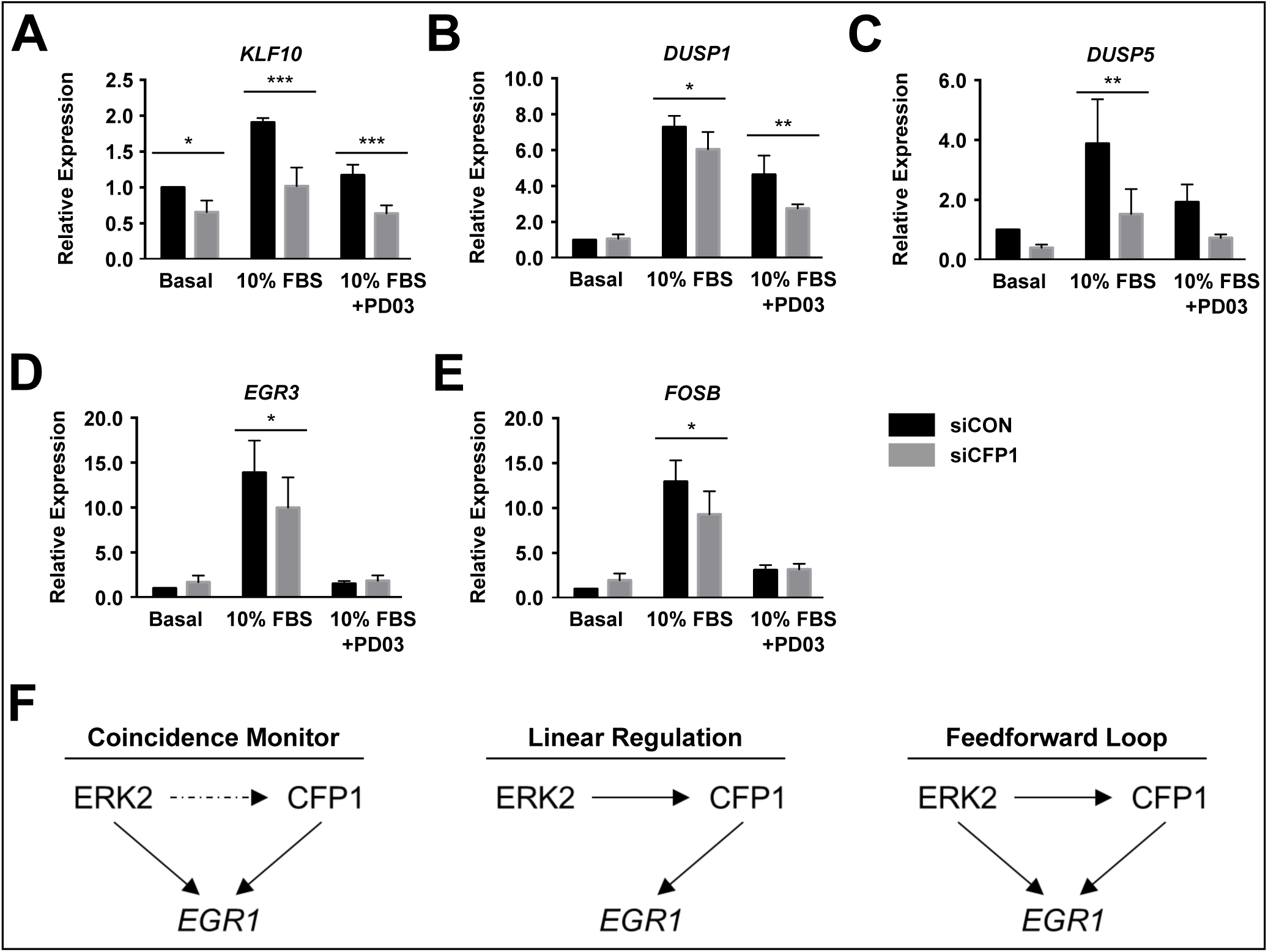
ERK1/2 signaling and CFP1 contribute differentially to serum-responsive regulation of target genes. **A-E** HeLa cells were subjected to 72 hours of siRNA treatment with either control (black bars) or CFP1-targeted (grey bars) oligonucleotides. Cells were serum-starved for 4 hours prior to pretreatment with 500 nM PD0325901 or vehicle control and subsequently stimulated with 10% FBS for 30 min. Total mRNA was extracted from cells, transcribed to cDNA and qRT-PCR was performed for the indicated target mRNAs. All changes were normalized to the siControl basal condition. Error bars represent standard deviation of three separate experiments. Statistical significance was assessed by Student’s two-tailed t-test: * = p<0.05; ** = p<0.005. **F** Models of non-mutually exclusive modes of regulation of serum responsive genes by ERK1/2 and CFP1.

### CFP1 is an ERK2 substrate in vitro and is phosphorylated on multiple sites

Because we were unable to detect direct CFP1-ERK2 binding in whole cell lysates without the presence of chromatin or formaldehyde cross-linking, we tested the possibility that transient interactions with ERK1/2 could result in CFP1 phosphorylation. Activated ERK2 phosphorylated full length recombinant CFP1 in an *in vitro* kinase assay with radiolabeled ATP (**Fig. 4A**), and subsequent phosphoamino acid analysis indicated CFP1 was modified on serine and threonine residues (**Fig. 4B**). The initial kinase assay results presented multiple bands of phosphorylated CFP1 (Fig. 4A, lane 3), suggesting multiple modified serine/threonine residues, and consistent with this, in longer assays with activated ERK2, a visible size shift consistent with multiple sites of phosphorylation on CFP1 was detected (**Fig. 4C**). CFP1 fragments incubated with activated ERK2 were also phosphorylated (**Fig. 4D**), and mass spectrometry analysis revealed the identities of several modified residues, including two sites that were predicted *in silico* [41], S224 and T227 (**Fig. 4E**). Ectopic expression of CFP1 point mutants at these sites did not result in major changes of global H3K4me3 when measured in whole cell lysate (**Fig. S4A**). Notably, Flag-CFP1-T227V, but not Flag-CFP1-S224A, was able to robustly transactivate a CpG-rich promoter-conjugated GFP construct relative to wildtype Flag-CFP1 (**Fig. S4B**). Combinatorial mutants also did not alter global H3K4me3 (**Fig. S4C**), but we cannot rule out compensatory post-translational modification given the extent of CFP1 phosphorylation by ERK2.

**Figure 4.**
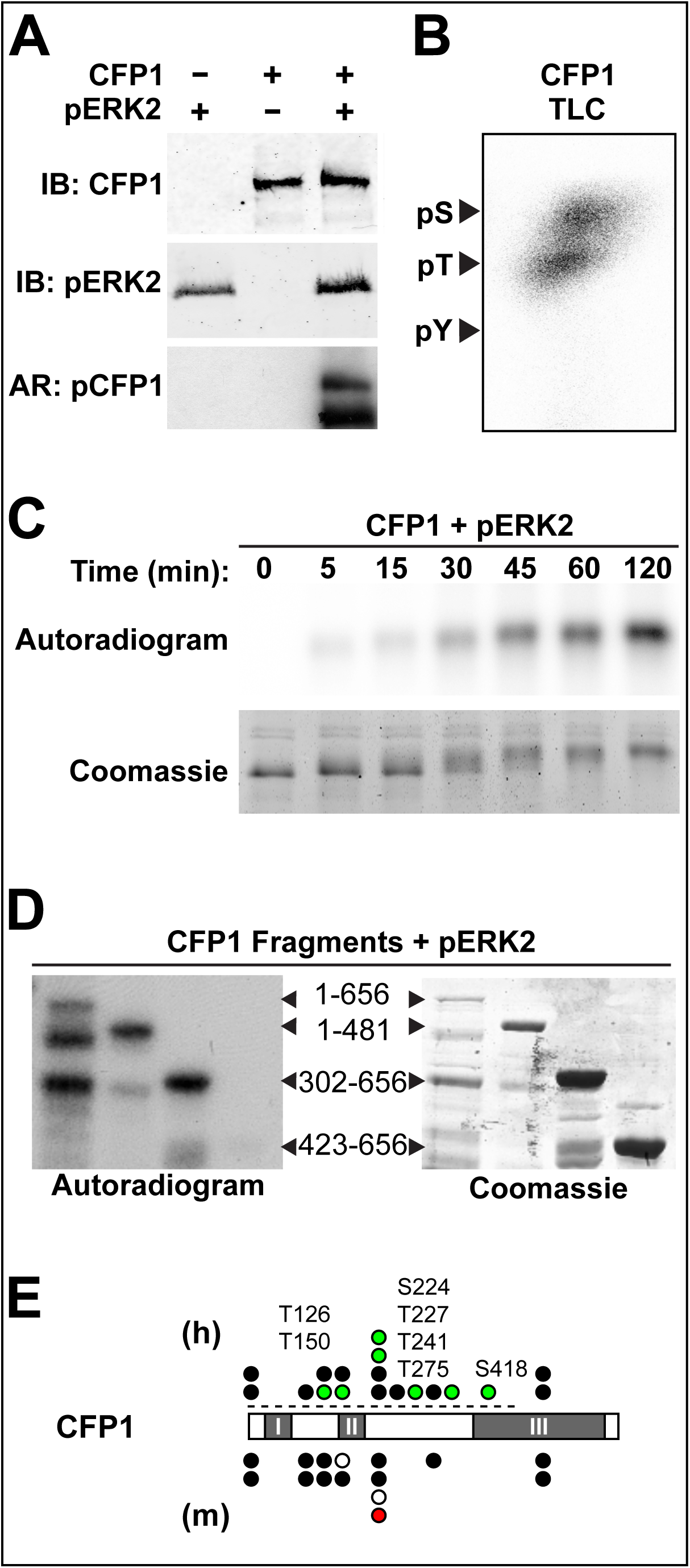
CFP1 is phosphorylated on multiple sites by activated ERK2 *in vitro*. **A** *In vitro* kinase reactions included full-length recombinant human CFP1 and activated recombinant ERK2 in the presence of [γ-^32^P]ATP at 30°C for 30 min. Proteins were resolved by SDS-PAGE and immunoblotted (IB) with indicated antibodies and subjected to autoradiography (AR). **B** Phosphorylated CFP1 from (A) was acid-hydrolyzed prior to separation of amino acid species by electrophoresis on a silica gel plate. Assignment of phosphorylated amino acids was made based on relative migration of pS/pT/pY standards, indicated by arrows. **C** Kinase reactions with activated recombinant ERK2 and a recombinant CFP1 fragment encompassing residues 1-481 proceeded for indicated amounts of time at 30°C. Results are representative of 3 independent experiments. **D** Full-length and fragments (indicated by arrows) of CFP1 subjected to *in vitro* kinase assay for 30 min at 30°C indicate multiple phosphorylation sites. Results are representative of 3 independent experiments. **E** CFP1 contains multiple phosphorylation sites identified by mass spectrometry and strongly predicted by previous reports and/or primary sequence. Black circles indicate all sites identified and banked on phosphosite.org [42], white circles indicate identified sites on S/T-P sequences, red circles indicate identified sites on P-x-S/T-P sequences, and green circles indicate sites identified by mass spectrometry (MS) in this study, with sites labeled by position above. Human (h) and mouse (m) CFP1 are highly conserved and both contain I. plant homeobox domain (PHD), II. CXXC domain, and III. coiled coiled (CC) domain. The region beginning shortly upstream of the CC domain and ending at the C-terminus was identified by yeast two-hybrid screen described in 1A. The dashed line indicates portion of hCFP1 subjected to MS analysis.

### Predicted ERK2 phosphorylation of SETD1B is confirmed in vitro

We postulated that ERK1/2-CFP1 co-regulation of target genes is likely to involve other catalytic factors that interact with CFP1. Comparisons of publicly available mass spectrometry data of DNMT1, SETD1A, and SETD1B reveal that all three contain multiple putative MAPK-targeted sites [42]. In particular, SETD1B displays the highest enrichment of prototypical MAPK recognition motifs S/T-P or P-X-S/T-P (**Fig. 5A**). CFP1 depletion results in no significant change in relative SETD1A mRNA expression (**Fig. 5B**), in contrast to a marked increase of relative SETD1B mRNA under the same conditions (**Fig. 5C**). This is consistent with a previous report that demonstrated SETD1A is regulated independently of SETD1B [43]. Intriguingly, CFP1 knockdown results in a notable increase in detected SETD1B protein, in contrast with the decrease in mRNA and suggestive of multiple levels of expression regulation (**Fig. 5C**). Finally, *in vitro* kinase assays with recombinant SETD1B (either with or without an intact GST affinity tag) confirm that both activated ERK1 (pERK1) and ERK2 can phosphorylate SETD1B (**Fig. 5D**).

**Figure 5.**
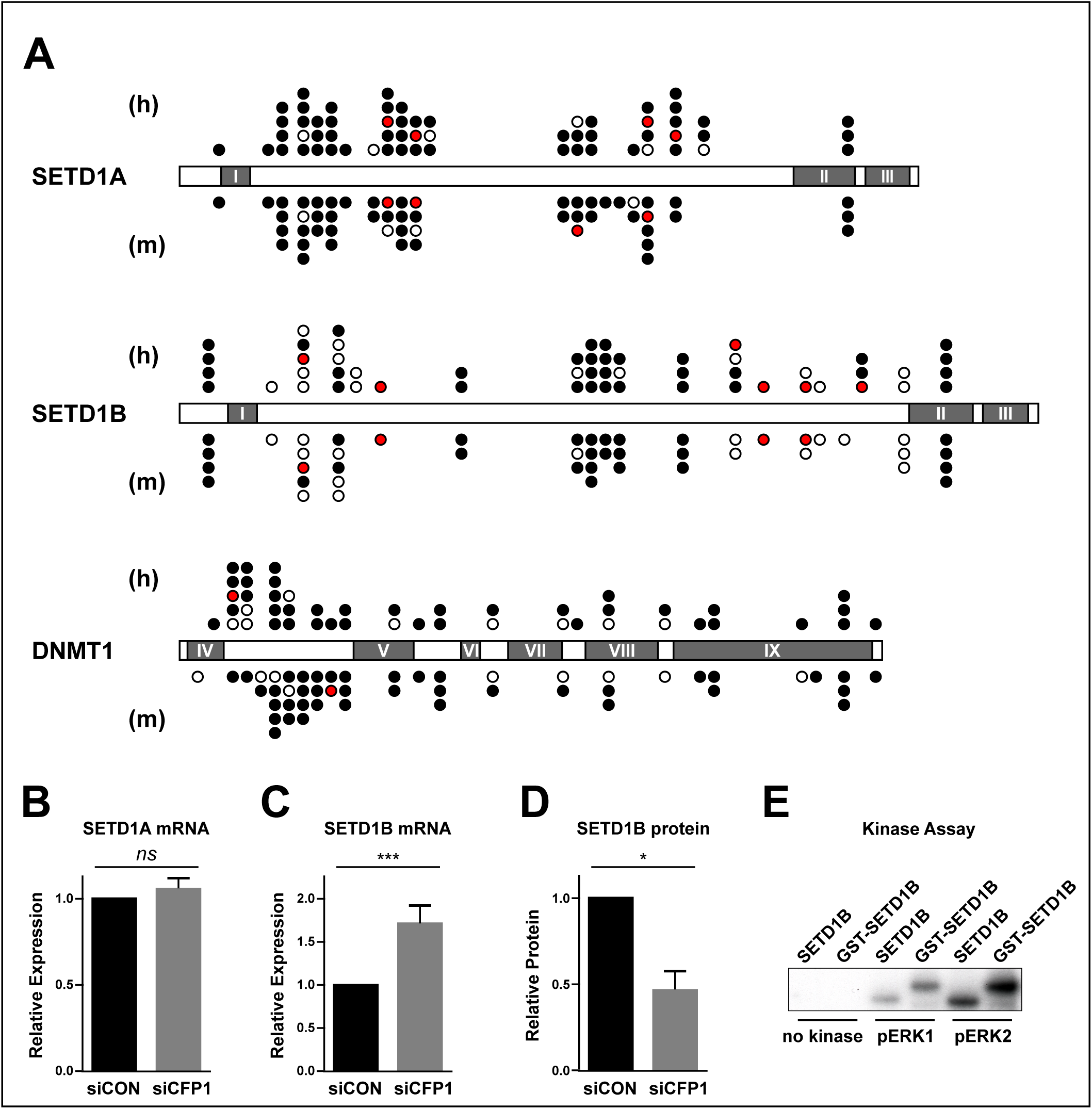
SETD1B, a CFP1-interacting protein, is also an ERK2 substrate. **A** SETD1A, SETD1B, and DNMT1 contain multiple phosphorylation sites. Notations are same as in Fig. 1E. SETD1A and SETD1B contain I. RNA recognition motifs, II. N-SET domains, and III. SET domains. DNMT1 contains IV. DMAP-binding domain, V. cytosine specific DNA methyltransferase replication foci domain, VI. CXXC zinc-finger domain, VII and VIII. Bromo-adjacent homology domains, and IX. DNA methyltransferase catalytic domain. **B** Relative expression of *SETD1A* mRNA measured upon siRNA-mediated depletion of CFP1. **C** Relative *SETD1B* mRNA was measured upon CFP1 depletion (*left*) along with SETD1B protein (*right*) under the same conditions. Results are representative of 3 independent experiments. **D** Radiolabeled kinase assay with recombinant SETD1B with or without a GST tag as substrate and activated ERK1 or ERK2 (pERK1 and pERK2, respectively).

## Discussion

ERK1 and ERK2 can mediate large-scale cellular dynamics and maintain homeostatic control by rapid response to upstream signals. Many of these signaling processes elicit regulated changes in gene expression, consistent with the abundance of evidence showing ERK1/2 form extensive interactions with chromatin and associated factors (reviewed in [6]). To better understand how ERK1/2 signaling influences genetic output, we sought to identify novel ERK2-interaction partners that are central to chromatin regulation. To this end, we utilized a yeast-two hybrid approach that employed selective activation of ERK2 to compare relative affinities for binding partners between phosphorylated and unphosphorylated states [34, 38, 44]. By this method we identified the epigenetic regulator CFP1 as a factor that interacts with ERK2 in eukaryotic cells.

The majority of literature on CFP1 is in a developmental signaling context, where it is known to be important for both DNA methylation and H3K4me3 deposition [39]. In particular, CFP1 has been most widely studied in mouse embryonic stem cells (mESCs), where ERK1/2 signaling is required for differentiation (in contrast to human ESCs, where ERK1/2 are critical for self-renewal) [45, 46]. ESCs derived from CFP1-knockout mice self-renewed but could not differentiate following removal of the cytokine leukemia inhibitory factor (LIF) from their culture medium [25, 47]. These cells exhibited a 70% global deficit in DNA methylation, with detectable hypomethylation at satellite repeats, stably methylated gene promoters and imprinted genes. Subsequent experiments suggested impairment of maintenance, but not *de novo* methyltransferase activity [25, 47]. Although both ERK1/2 signaling and CFP1 are known to regulate DNMT1 expression [22-24, 48], depletion of CFP1 or inhibition of ERK1/2 signaling did not result in a global reduction of 5mC in HeLa cells, a cancer cell line.

In addition to interacting with DNMT1 and regulating DNA methylation, CFP1 is also a member of two histone methyltransferase complexes, SETD1A and SETD1B. Both of these complexes mediate H3K4me3 at target genes and associate with largely non-overlapping regions of chromatin [49]. Notably, chromatin immunoprecipitation coupled to deep sequencing (ChIP-seq) studies indicate extensive colocalization of ERK1/2 and H3K4me3 [11]. Loss of CFP1 in mESCs resulted in aberrant placement of H3K4me3 to chromatin regions outside promoters, indicating CFP1 is essential for proper targeting of SETD1A/B complexes. Previous work has shown that SETD1A and SETD1B are subject to different upstream signals, suggesting that CFP1 regulation is highly context-dependent [43, 49]. Supporting this notion, CFP1 has also been shown to mediate rapid H3K4me3 enrichment in a gene-induction response in B-cells [18], however, no such role for SETD1B has been reported to date. Our work demonstrates that both CFP1 and SETD1B are ERK2 substrates and suggests that interactions with these factors modulate ERK1/2 signaling functions in the nucleus. In this regard, we selected three positively-driven regulation modes to highlight (coincidence monitoring, linear regulation, and a feedforward loop) because any of these non-mutually exclusive models may explain our expression induction results.

Much like their wide-ranging roles in regulation of cytosolic substrates, ERK1/2 signaling in the nucleus is abundant if not well understood. As previously discussed, ERK1/2 have evolved divergent roles in development between humans and mice, suggesting adaptability is inherent to ERK1/2 biochemical activity. In addition to varied roles in development, ERK1/2 are required for a diversity of signal-dependent responses, further suggesting that vulnerability to misregulation is a consequence of its versatility.

As the effector kinases of the Ras-Raf-MEK-ERK pathway, ERK1/2 are crucial for normal cellular function and prone to driving disease states under conditions of inappropriate signaling. An improved understanding of ERK1/2 influence on gene expression is critical for eventual development of more finely tuned treatment options. Here, we describe identification of key substrates of the ERK1/2 pathway to clarify how one transduction relay is capable of a diverse array of outputs.

## Conclusions

Our work concludes that ERK2 and CFP1 are both critical for optimal serum-responsive gene induction, that ERK2 and CFP1 interact on chromatin and further that CFP1 is an ERK2 substrate *in vitro*. The H3K4me3 deposition factor SETD1B, known to interact with CFP1, is also an ERK2 substrate *in vitro*.

## Methods

### Cell culture and transfections

HeLa cells were obtained from ATCC and cultured in Dulbecco’s Modified Eagle Medium (DMEM) supplemented with 10% v/v fetal bovine serum (Sigma) and 2 mM L-glutamine. Human CFP1 cDNA subcloned into p3xFLAG-CMV-7.1 (Sigma E4023) and point mutations generated using site-directed mutagenesis were transfected with Lipofectamine 2000 (Life Technologies). All constructs were Sanger-sequenced to verify that lack of spurious mutations. Duplex siRNA oligonucleotides targeting human CFP1 (s26937) or a nontargeting control (4390843) were acquired from Life Technologies. Oligonucleotides were reverse transfected into HeLa cells at 20 nM for 72 hours with RNAiMax (Life Technologies) according to the manufacturer’s instructions. For Flag-ERK2 immunoprecipitations, cells were transfected with p3xFLAG-CMV-ERK2 [29] for 48 hours with Lipofectamine 2000.

### Cell harvest

For whole cell lysates, cells were lysed with RIPA buffer (50 mM Tris pH7.4, 150 mM NaCl, 1 mM EDTA, 1% NP-40, 0.5% sodium deoxycholate, 0.1% SDS, 80 mM beta-glycerophosphate, 100 mM NaF and 2 μM Na_3_VO_4_) with 1:1000 protease inhibitor cocktail (in 62.5 mL DMSO: 25 mg pepstatin A, 25 mg leupeptin, 250 g N_a_-p-tosyl-L-arginine methyl ester, 250 mg N_a_-p-tosyl-L-lysine chloromethyl ketone hydrochloride, 250 mg N_a_-Benzoyl-L-arginine methyl ester and 250 mg soybean trypsin inhibitor), and sonicated prior to immunoblotting.

### Nuclear fractionation

Cells were rinsed with phosphate-buffered saline (PBS) and resuspended at 10^7^ cells per mL in 10 mM HEPES (pH 7.9), 10 mM KCl, 1.5 mM MgCl_2_, 0.34 M sucrose, 10% glycerol, 1 mM dithiothreitol (DTT). Triton X-100 was added to a final concentration of 0.1% and cells were incubated on ice for 5 minutes. Crude nuclear fractions were pelleted in a microfuge at 3500 rpm for 5 min. For further isolation of chromatin, nuclear pellets were washed once in the buffer above without Triton then resuspended in 3 mM EDTA, 0.2 mM EGTA, 1 mM DTT for 15 min on ice with periodic vortexing. Chromatin was pelleted in a microfuge at 4000 rpm for 5 min.

### Mononucleosome preparation and immunoprecipitation

Mononucleosomes were prepared by resuspending chromatin pellets with Buffer A (10 mM HEPES pH 7.9, 10 mM KCl, 1.5 mM MgCl_2_, 0.34 M sucrose, 1 mM DTT, 10% glycerol) containing 1 mM CaCl_2_, then digested at room temperature with micrococcal nuclease (Thermo Scientific). Digestions were stopped by adding EGTA to a final concentration 1 mM. Undigested chromatin was pelleted by centrifugation at 4000 rpm for 5 min. Supernatants were used for immunoprecipitation with the indicated antibodies as previously reported [30]. For immunoprecipitation, salt concentrations of the mononucleosome fraction were adjusted to 150 mM with KCl and Triton X-100 was added to 0.2%. Flag-ERK2 was immunoprecipitated using Flag M2-conjugated agarose (Sigma) and washed at least three times with salt-adjusted Buffer A prior to denaturation by boiling at 100°C in Laemmli sample buffer.

### Acid-extraction of histones

Cells were harvested in ice cold 1X PBS and resuspended in cold hypotonic lysis buffer (10 mM Tris pH 8.0, 1 mM KCl, 1.5 mM MgCl2, 1 mM DTT) prior to acid extractions of histones as described [31]. Total histone content was measured after separation by 15% SDS-PAGE and Coomassie staining with protein standards.

### Immunoblotting and LiCor imaging

Proteins were separated by SDS-PAGE and transferred to nitrocellulose membranes (Millipore). Membranes were blocked with LiCor blocking buffer and incubated with the indicated antibodies. Fluorescently labeled secondary antibodies were used in conjunction with the LI-COR Odyssey dual-colour system, and band quantitation was performed using Image Studio software. Western blot analysis was performed using the following primary antibodies: rabbit anti-CFP1 (Bethyl Laboratories); rabbit anti-ERK1/2 (produced in-house [32]); mouse anti-pERK1/2 (T185/Y187) and mouse anti-Flag M2 (Sigma); and mouse anti-H3 and rabbit anti-H3K4 trimethyl (Active Motif).

### DNA Methylation assay

Genomic DNA was isolated with *Quick*-DNA™ Miniprep Plus Kit (Zymo Research) and global 5mC content was measured by MethylFlash Methylated DNA 5-mC Quantification Kit (Epigentek) according to manufacturer’s instructions.

### Microarray and qRT-PCR

Total RNA was prepared from HeLa cells using the Pure-link RNA Mini Kit (Life Technologies) according to the manufacturer’s instructions. Briefly, RNA was checked for concentration and quality using an Agilent 2100 Bioanalyzer, then cRNA was synthesized and labeled prior to hybridization to an Affymetrix Human Transcriptiome 2.0 Array chip and detection. Raw data was analyzed using Affymetrix Transcriptome Analysis Console 2.0 software. For qRT-PCR, cellular RNA was isolated with TRI reagent (Applied Biosystems) and reverse transcribed using the iScript cDNA synthesis kit (BioRad). qRT-PCR was performed with SYBR Green Supermix (BioRad) and fluorescence was measured using a quantitative real-time thermocycler (BioRad). Relative changes in gene expression calculated by 2^−ΔCt^, with *actin* as internal expression control.

### Recombinant protein expression and purification

His_6_-tagged ERK2 was expressed in Origami *E. coli* (EMD Millipore) using the NpT7-His_6_-ERK2 construct as described previously [33]. ERK2 was activated in bacteria by coexpressing ERK2 with MEK1 R4F [34] as described [35]. Human CFP1 and truncation constructs were subcloned into the pGEX-6P-1 vector encoding an N-terminal glutathione S-transferase (GST) tag (GE Healthcare) prior to expression in Origami cells. Fusion proteins were induced overnight at 20°C with 0.4 mM isopropyl-1-thio-β-D-galactopyranoside (IPTG). Cells were lysed in buffer containing 20 mM HEPES (pH 7.5), 1 mM EDTA, 1 mM DTT with 1:500 protease inhibitor cocktail (described above in 2.2). Pellets were lysed and clarified, and GST-tagged proteins were bound to glutathione resin according to the manufacturer’s instructions (Pierce). Protein-bound resin was washed in lysis buffer at least three times and bound protein was eluted with 20 mM glutathione. GST tags were cleaved with PreScission protease per the manufacturer’s instructions (GE Healthcare). Isolated CFP1 fragments were dialyzed in 25% glycerol, 150 mM NaCl, 10 μM ZnCl_2_, 1 mM EDTA and 1 mM DTT.

### In vitro kinase assay and mass spectrometry

In vitro kinase reactions were carried out at 30°C in the presence of 10 mM Tris (pH 7.5), 10 mM MgCl_2_, 200 μM ATP and 10 μCi [γ-^32^P]ATP. Reactions were quenched by addition of 5x Laemmli sample buffer and boiled for 2 min at 100°C. Samples for mass spectrometry were phosphorylated for 60 min without radiolabeled ATP, separated by SDS-PAGE, stained with Coomassie blue and digested with either trypsin or elastase to maximize protein coverage. Sites were called using the ModLS algorithm, using cutoff values for positive site identification that represent a scenario where the false discovery rate < 1% [36].

### Phosphoamino acid analysis

Full-length recombinant CFP1 was phosphorylated in vitro with pERK2 and [γ-^32^P] ATP for 30 minutes. Phospho-CFP1 was immunoblotted and exposed to film to verify phosphate incorporation, then the band was excised and hydrolyzed for 1 hr in 6 N HCl. Hydrolysis products were resolved by thin layer electrophoresis as described [37]. The plate was exposed to film, and identities of phosphorylated residues were deduced by co-migration of phosphoamino acid standards visualized with ninhydrin.

## Abbreviations

ERK: extracellular signal-regulated kinase
CFP1: CXXC-finger protein 1
H3K4me3: histone H3 lysine 4 trimethylation
5mC: 5-methylcytosine (DNA methylation)
IEG: immediate early gene
EGR: early growth response
DUSP: dual specificity phosphatase
FOSB: FBJ murine osteosarcoma viral oncogene homolog B
ESC: embryonic stem cell
SET: Su(var)3-9, Enhancer-of-zeste and Trithorax
DNMT: DNA methyl-transferase.

## Declarations

### Ethics approval and consent to participate

Not applicable

### Consent for publication

Not applicable

### Availability of data and material

All data generated or analyzed during this study are included in this published article (and its supplementary information files).

### Authors’ contributions

M.H.C. and H.M. provided guidance and important intellectual content for the report and made substantial contributions to data interpretation. A.S.K, A.M.K, and M.H.C. drafted the manuscript and revised it critically based on contributions from all authors. A.S.K. and A.M.K. made substantial contributions to conception and design of experiments, acquisition of data, data analysis, and interpretation of data. S.E., S.S, K.M, and D.C.T. made substantial contributions to acquisition of data and analysis, and D.C.T. and C.W. were instrumental for data analysis and interpretation.

## Acknowledgements

The authors thank Drs. Michael Kalwat and Clinton Taylor, along with other current and former members of the Cobb laboratory for valuable discussions, and Dionne Ware for administrative assistance.

## Supplementary Figure Legends

**Figure S1. Subcellular fractionation of HeLa cells following stimulation and inhibition of ERK1/2.** HeLa cells were starved to quiescence in serum-free DMEM, then pretreated with 500 nM PD0325901 or DMSO for one hour prior to stimulation. EGF was added for 15 minutes at 10 ng/mL. Cells were fractionated to separate crude cytosolic, soluble nuclear and insoluble chromatin pools, and fractions were volume-normalized prior to separation by SDS-PAGE and immunoblotting with the indicated antibodies. Antibodies against tubulin and histone H3 were used to validate the quality and consistency of cytosolic and nuclear fractions, respectively.

**Figure S2. CFP1 depletion by siRNA.** Cells were transfected with either siRNA against CFP1 (gene name *CXXC1*) or a control target for 72 hours, then incubated in serum-free DMEM prior to a 15 minute treatment with 10% FBS. Five minutes before stimulation, cells were treated with either 500 nM PD0325901 or DMSO. Cells were then lysed and proteins resolved by SDS-PAGE prior to blotting with the indicated antibodies.

**Figure S3. Fold induction of ERK1/2-regulated genes with and without CFP1 depletion.** HeLa cells were subjected to 72 hours of siRNA treatment with either control or CFP1-targeted oligonucleotides. Cells were lysed with Trizol and mRNA was extracted. qRT-PCR was performed for the indicated target mRNAs. All changes were normalized to the knockdown-matched basal condition. Error bars represent standard deviation of three separate experiments. Statistical significance was assessed on standard errors by Student’s two-tailed t-test; ** = p < 0.05 and *** = p <0.005.

**Figure S4. Characterization of CFP1 point mutants. A** Overexpression of CFP1 and mutants does not significantly alter global H3K4me3 as measured from whole cell lysate. **B** HeLa cells transfected with equivalent amount of pEGFP plasmid and a range of Flag-CFP1. Signal intensity of GFP normalized to actin, and relative modification normalized to total actin was plotted. Error bars indicate standard deviation of three separate experiments. Significance was quantified based on standard errors by Student’s two-tailed t-test; * = p < 0.05. **C** Expression of combinatorial mutants (EV, empty vector control; CFP1 3xM, S124A S224A T227V; CFP1 4xM, S124A T126V S224A T227V) of CFP1 do not alter global H3K4me3 as measured in whole cell lysates.

