## Supplementary figures and images for "ERK2 phosphorylates the epigenetic regulator CXXC-finger protein 1 (CFP1)"

### Figure S4

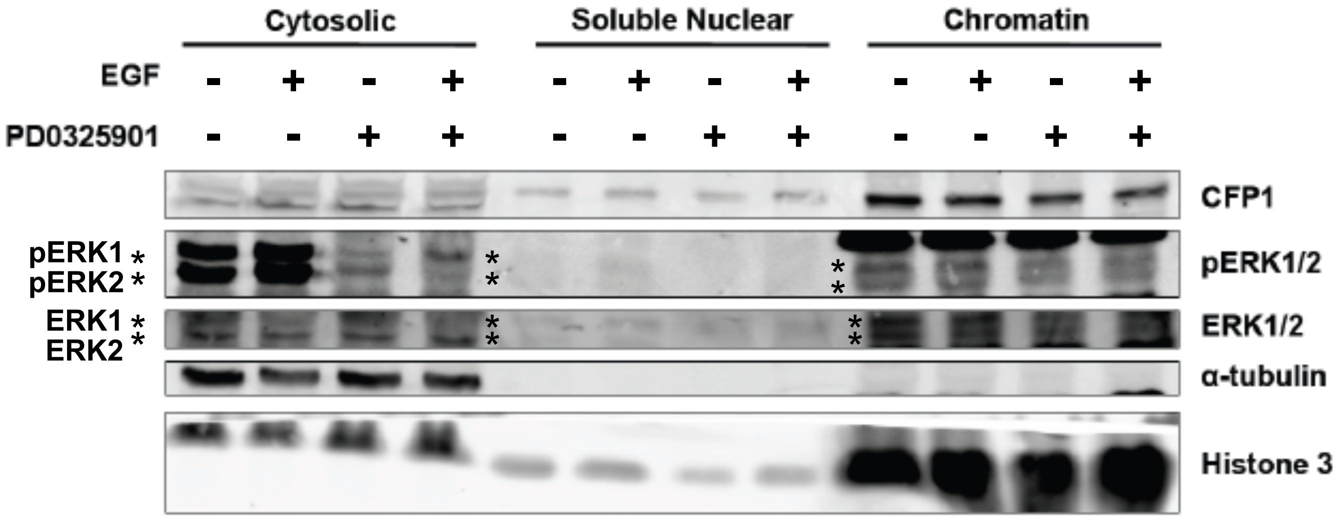


#### Supplementary Figure 1

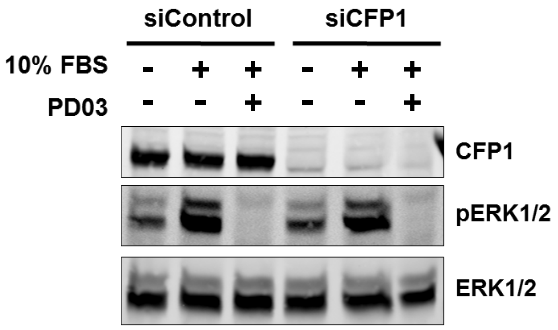


#### Supplementary Figure 2

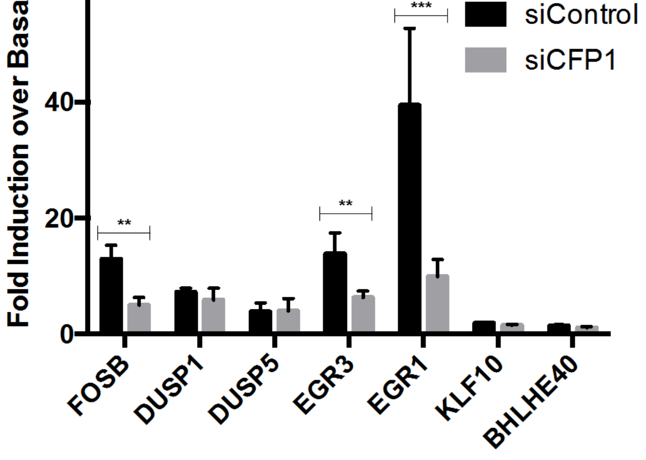


#### Supplementary Figure 3

### Figures S1-S3

A

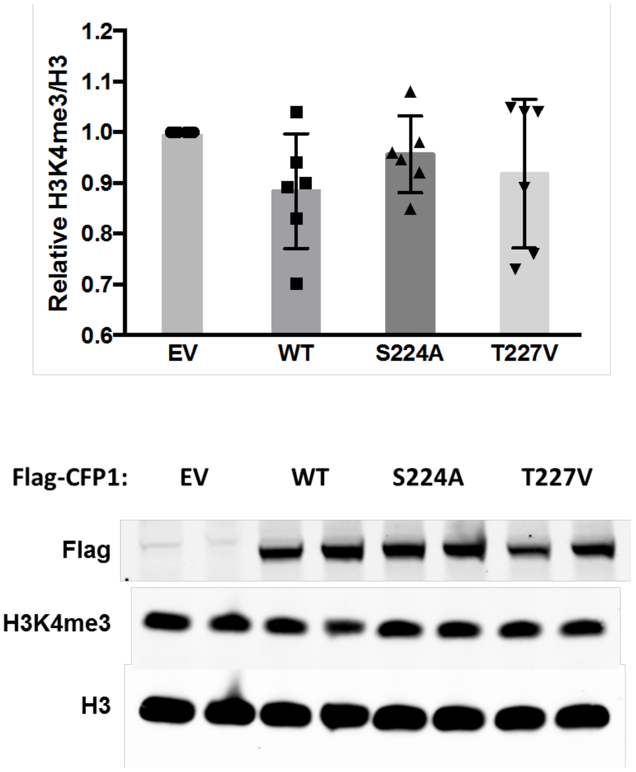


B

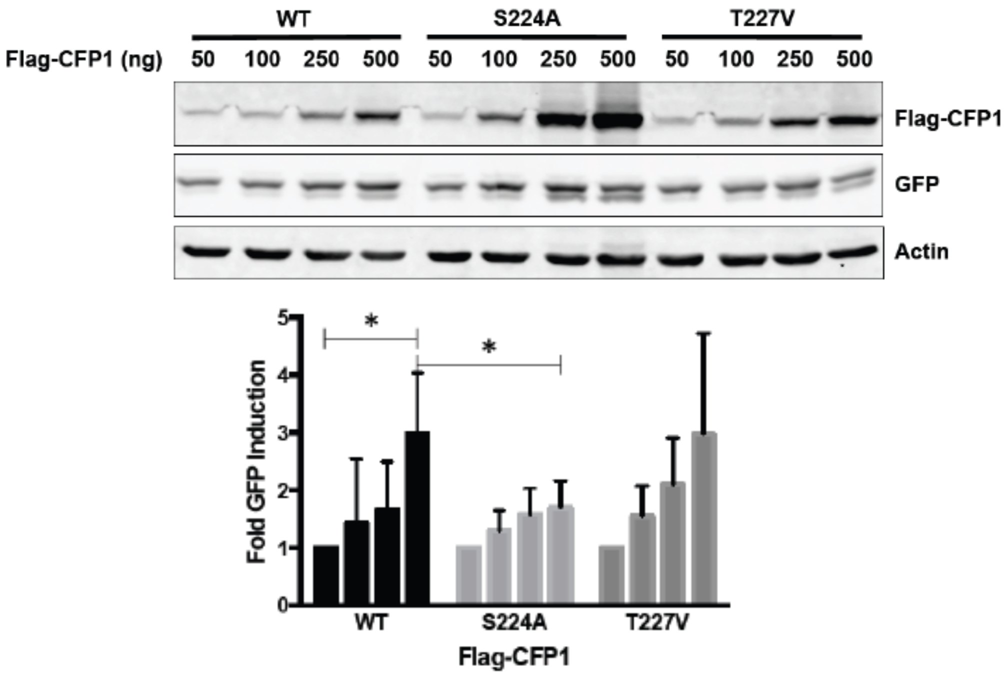


C

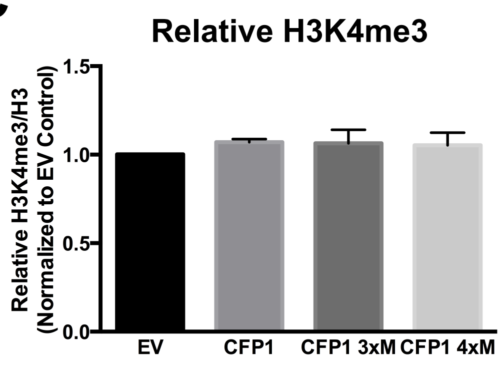


#### Supplementary Figure 4
